# National Institutes of Health Institute and Center Award Rates and Funding Disparities

**DOI:** 10.1101/2020.12.27.424490

**Authors:** Michael Lauer, Jamie Doyle, Joy Wang, Deepshikha Roychowdhury

## Abstract

A previous report found an association of topic choice with race-based funding disparities among R01 applications submitted to the National Institutes of Health (“NIH”) between 2011-2015. Applications submitted by African American or Black (“AAB”) Principal Investigators (“PIs”) skewed toward a small number of topics that were less likely to be funded (or “awarded”). It was suggested that lower award rates may be related to biases of peer reviewers. However, the report did not account for differential funding ecologies among NIH Institutes and Centers (“ICs”). In a re-analysis, we find that 10% of 148 topics account for 50% of applications submitted by AAB PIs. These applications on “AAB Preferred” topics were funded at lower rates, but peer review outcomes were similar. The lower rate of funding was primarily due to their assignment to ICs with lower award rates. After accounting for IC-specific award rates, topic choice was not associated with funding.

## Introduction

Data recently reported by Hoppe et al [Hoppe et al., 2019] from the National Institutes of Health (“NIH”) suggest that part of the well-documented funding disparity [Ginther et al., 2011] affecting African-American Black (“AAB”) principal investigators (“PIs”) may be related to the topic of their applications. The authors of that report (including the first author of this report) found that topic choice accounted for over 20% of the disparity and wondered whether biases on the part of peer reviewers might explain why some application topics fare less well when submitted to the NIH for consideration of funding.

However, peer review outcomes are not the only determinant of funding. Applications submitted to the NIH are assigned to one of 24 grant-issuing institutes or centers (“ICs”) that in turn make decisions about which proposal to fund. The proportion of applications funded (or “award rate”) varies accross ICs; therefore, we can think of the NIH process as not one competition hinging entirely on peer review but rather 24 separate competitions. The variability of award rates relates to differences in number of applications each IC receives, available funds, and IC priorities.

Hoppe et al [Hoppe et al., 2019] did not account for IC assignment or variation in IC-specific award rates. It is possible that the apparent link between topic choice and funding disparities may reflect differences in IC assignment, since ICs receive applications according to alignment with their stated mission. For example, applications focusing on cancer epidemiology are more likely to be assigned to the National Cancer Institute while those focusing on basic human biology are more likely to be assigned to the National Institute of General Medical Sciences. If award rates at the National Institutes of General Medical Sciences are higher than at the National Cancer Institute, it might appear that NIH “prefers” basic human biology over cancer epidemiology. While the former topic does fare better with a higher likelihood of funding, this may be largely because of different IC award rates as opposed to differences in how the topics are received by peer reviewers.

We therefore re-analyzed the data from Hoppe et al [Hoppe et al., 2019] focusing on the possible role of IC assignment in application outcomes. To minimize biases related to repeated looks by the peer system on individual proposals (from resubmissions [Lauer, 2017] or competing renewals [Lauer, 2016]) we focus on *de novo* applications submitted to the NIH for the first time.

## Methods

These analyses are based on R01 applications submitted to NIH between 2011 and 2015. Hoppe et al [Hoppe et al., 2019] described in detail NIH peer review processes and the “Word2vec” algorithm [Mikolov et al., 2013] used to designate a topic for each application. Briefly, each application is assigned to a peer review group. After a preliminary pre-meeting review, approximately half are deemed to be potentially meritorious and are therefore discussed during a formally convened meeting. After the meeting, each discussed application receives a priority score ranging from 10 (best) to 90 (worst); many, but not all, applications also receive a “percentile ranking” to account for differences in how individual peer review groups calibrate their scores.

Applications are not only assigned to peer review groups; they are also assigned to ICs. ICs ultimately make decisions about which applications to fund, with funding decisions based on peer review scores, strategic priorities, and availability of funds.

To eliminate biases due to prior reviews, we focused on applications coming to the NIH for the first time; in other words, we excluded resubmissions [Lauer, 2017] and competing renewals [Lauer, 2016]. For each IC, we calculated award rates as number of applications funded divided by number of applications assigned. We also noted what proportion of applications had a principal investigator (“PI”) who self-identified as AAB. For multi-PI applications, only the demographic information of the contact PI was used. We designate those ICs in the top quartile of AAB application proportions as “Higher AAB” ICs.

There were 148 topics identified by the Word2vec algorithm [Mikolov et al., 2013]. For each topic, we counted the number of applications of submitted by AAB PIs. Consistent with the findings of Hoppe topics were not randomly distributed by PI race; there were 15 topics that accounted for 50% of applications submitted by AAB PIs. We designate these applications as having “AAB Preferred” topics.

To assess the association of topic choice, IC assignment, and peer review on application success, we compared peer review and funding outcomes according to whether applications were assigned to Higher AAB ICs and separately whether applications topics were AAB Preferred or Other. We performed a series of probit regression analyses with funding as the dependent variable and AAB PI as an explanatory variable. We added topic choice (AAB Preferred or Other), IC assignment (Higher or Lower AAB), and IC award rate in separate models and examined whether the regression coefficient relating AAB PI to funding decreased, and if so, by how much. We conducted a separated set of probit analyses focusing on topic choice, either alone or after adjustment for IC-specific award rate and other possible confounders. Akaike Information Criteria, Bayesian Information Criteria, and Log Likelihood values informed model strength.

To assess the the association of topic choice with peer review outcomes, we focused on applications that were discussed and therefore received a priority score. We constructed a plot of topic-specific mean peer review scores according to number of applications in each topic. We expected to find a greater variance of mean scores for topics receiving fewer applications (“regression to the mean”). We generated a linear regression model to estimate a predicted mean score for each topic based on topic size, and calculated a residual for each topic by substracting from each topic-specific mean score the model-based predicted mean score.

All analyses used R [Rpr] packages, including tidyverse [Wickham et al., 2019], ggplot2 [Wickham, 2016], finalfit [fin], and texreg [Leifeld, 2013].

## Results

Of 157,405 applications received, there were, after exclusion of resubmissions and competing renewals, 99,195 applications considered by NIH for the first time. Of these 8422 were funded, for an overall award rate of 8%. There were 1685 applications, or 2%, submitted by AAB PIs. Table 1 shows IC-specific values for applications received, applications funded, award rates, and percent applications coming from AAB PIs. Of note, award rates varied from 6% to 15%, while the proportion of applications with AAB PIs ranged from <1% to nearly 15%.

### Review and Funding Outcomes According to IC and to Topic

Table 2 shows review and funding outcomes for applications according to whether the assignment was to an IC in the top quartile of AAB applications (“Higher AAB”). These ICs were the National Institute of Allergy and Infectious Diseases, the National Institute of Environmental Health Sciences, the National Institute of Child Health and Human Development, the National Institute of Minority Health and Disparities, the National Institute of Nursing Research, and the Fogarty International Center. Applications submitted to Higher AAB ICs were 3 times more likely to come from AAB PIs. Review outcomes – proportion discussed and, for those applications that were discussed at peer review meetings, priority scores and percentile rankings – were similar in both groups. Despite the similar review outcomes, they were 13% less likely to be funded.

Table 3 shows corresponding values according to whether applications were focused on the 15 topics that made up 50% of all applications with AAB PIs (“AAB Preferred” topics). Again, peer review outcomes were similar in the two groups, but applications focusing on AAB Preferred topics were 8% less likely to be funded.

**Table 1:**
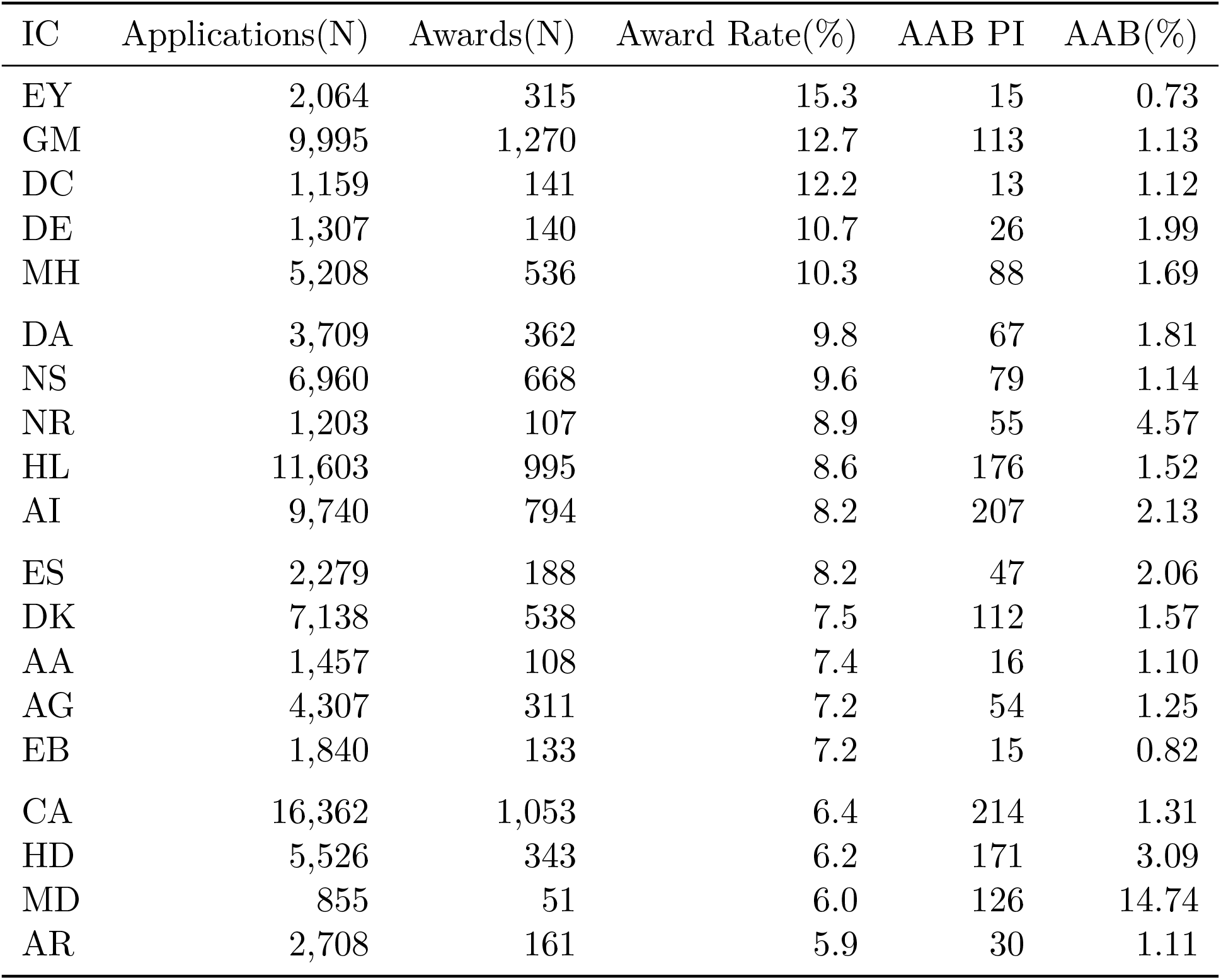
Application characteristics according to Institute or Center (IC). AAB = African American or Black; PI = Principal Investigator; EY = National Eye Institute; DC = National Institute of Deafness and Communications Disorders; GM = National Institute of General Medical Sciences; DE = National Institute of Dental and Craniofacial Research; MH = National Institute of Mental Health; DA = National Institute on Drug Abuse; NS = National Institute of Neurological Disorders and Stroke; NINR = National Institute of Nursing Research; HL = National Heart, Lung, and Blood Institute; AI = National Institute of Allergy and Infectious Diseases; ES = National Institute of Environmental Health Sciences; DK = National Institute of Diabetes and Digestive and Kidney Disease; AA = National Institute on Alcohol Abuse and Alcoholism; AG = National Institute on Aging; EB = National Institute of Biomedical Imaging and Bioengineering; CA = National Cancer Institute; HD = Eunice Kennedy Shriver National Institute of Child Health and Human Development; MD = National Institute on Minority Health and Health Disparities; AR = National Institute of Arthritis and Musculoskeletal and Skin Diseases. Data for ICs with cell sizes not exceeding 11 are not shown due to privacy concerns.

**Table 2:** Application review and funding outcomes according to whether Institute or Center received a higher or lower proportion of applications from AAB principal investigators. AAB = African American or Black; PI = Principal Investigator.

| Characteristic or Outcome |  | Higher AAB | Lower AAB |
| --- | --- | --- | --- |
| Total N (%) |  | 19701 (19.9) | 79494 (80.1) |
| PI AAB | Yes | 617 (3.1) | 1068 (1.3) |
| Discussed | Yes | 9090 (46.1) | 35608 (44.8) |
| Priority Score | Median (IQR) | 39.0 (30.0 to 48.0) | 40.0 (30.0 to 48.0) |
| Percentile Ranking | Median (IQR) | 33.0 (21.0 to 44.0) | 33.0 (21.0 to 44.0) |
| Awarded | Yes | 1494 (7.6) | 6928 (8.7) |

**Table 3:** Application review and funding outcomes according to whether topic was among those that accounted for the majority of AAB applications. Abbreviations as in Table 2.

| Characteristic or Outcome |  | AAB Preferred | Other |
| --- | --- | --- | --- |
| Total N (%) |  | 25013 (25.2) | 74182 (74.8) |
| PI AAB | Yes | 842 (3.4) | 843 (1.1) |
| Discussed | Yes | 11522 (46.1) | 33176 (44.7) |
| Priority Score | Median (IQR) | 40.0 (31.0 to 48.0) | 39.0 (30.0 to 48.0) |
| Percentile Ranking | Median (IQR) | 34.0 (22.0 to 45.0) | 33.0 (20.0 to 44.0) |
| Assigned IC AAB Proportion | Higher AAB | 6331 (25.3) | 13370 (18.0) |
| Awarded | Yes | 1999 (8.0) | 6423 (8.7) |

Why do applications on AAB Preferred topics have worse funding outcomes despite similar peer review assessments? Table 3 shows that applications on AAB Preferred topics are 41% more likely to be assigned to Higher AAB ICs. The scatter plot in Figure 1 shows IC award rate according to the proportion of applications assigned to it that focus on AAB Preferred topics. ICs with receiving a higher percentage of AAB Preferred topic applications have lower award rates.

**Figure 1:**
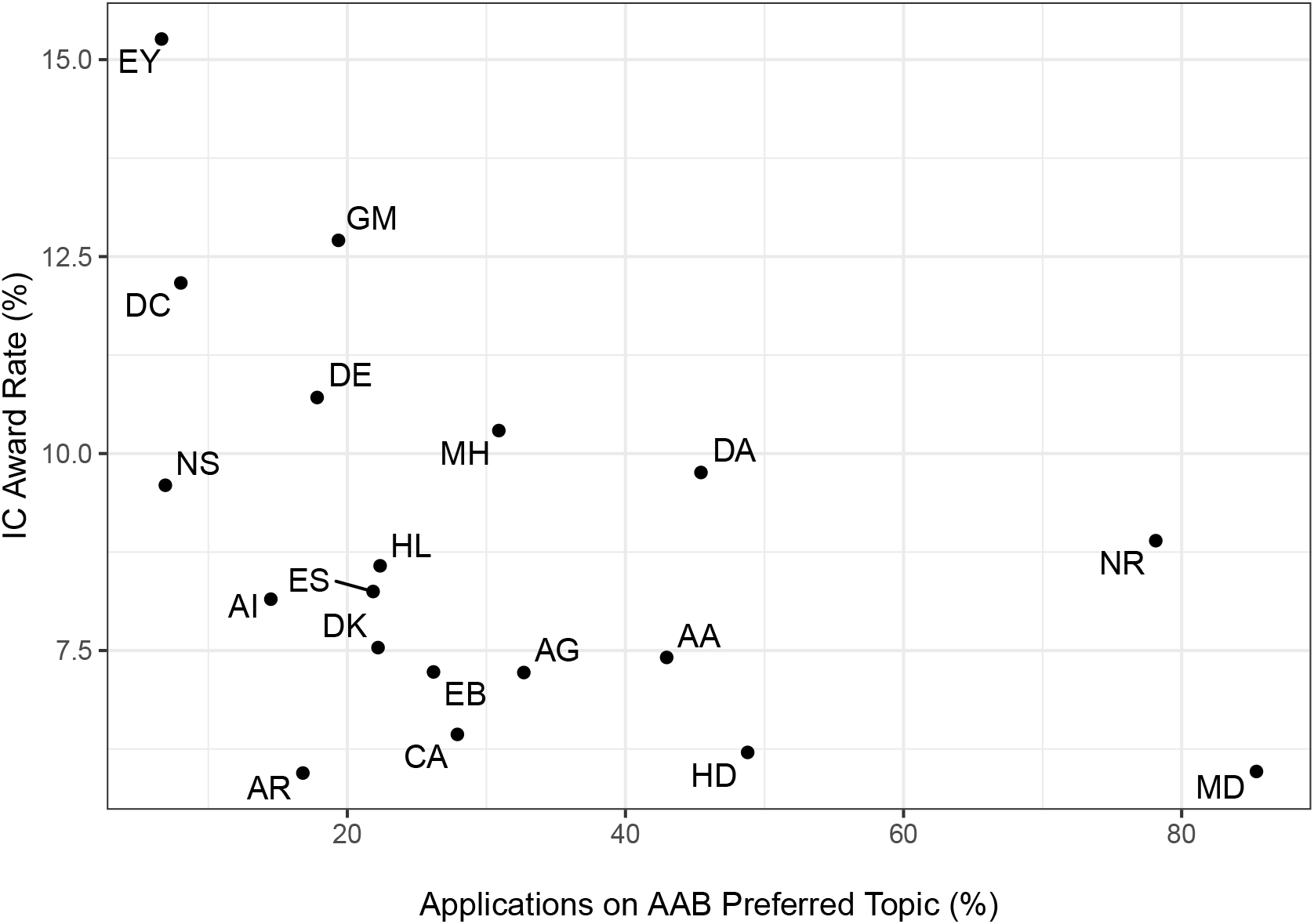
Scatter plot of IC specific award rates according to proportion of applications focusing on AAB Preferred topics. Abbreviations are the same as in Tables 1 and 2. ICs that receive relatively more applications on AAB Preferred topics have lower award rates. Data for ICs with cell sizes not exceeding 11 are not shown due to privacy concerns.

### Probit Regression Models

Table 4 shows the association of an application with an AAB PI with the probability of funding. Consistent with Hoppe et al [Hoppe et al., 2019] and prior literature [Ginther et al., 2011], AAB PI applications had a lower likelihood of funding (negative regression coefficient for AAB Principal Investigator). Adjusting for the topic (AAB Preferred or Other) reduced the regression coefficient for race by 5%; similarly adjusting for IC assignment (Higher or Lower AAB) reduced the regression coefficient by 6%. However, adjusting for the award rate of the assigned IC reduced the regression coefficient for race by 14%.

**Table 4:** Probit Regression Models (regression coefficients and standard errors) with focus on the PI. Model 1 shows the univariable association of funding success according to whether the PI is AAB. Model 2 adjusts for topic, Model 3 adjusts for IC assignment, and Model 4 adjusts for IC award rate. Note that the absolute value for the regression coefficient linking AAB PI to funding outcome decreases with each of these adjustments, with the greatest reduction after adjusting for IC award rate. AIC = Akaike Information Criterion; BIC = Bayesian Information Criterion; Num. obs. = Number of Observations. Other abbreviations same as in Tables 1 and 2.

|  | Model 1 | Model 2 | Model 3 | Model 4 |
| --- | --- | --- | --- | --- |
| Intercept | -1.370***<br>(0.006) | -1.360***<br>(0.007) | -1.356***<br>(0.006) | -1.910***<br>(0.020) |
| AAB Principal Investigator | -0.185***<br>(0.049) | -0.175***<br>(0.049) | -0.173***<br>(0.049) | -0.159**<br>(0.049) |
| AAB Preferred Topic |  | -0.040**<br>(0.013) |  |  |
| IC Higher AAB Applications |  |  | -0.072***<br>(0.015) |  |
| IC Award Rate |  |  |  | 0.062***<br>(0.002) |
| AIC | 57637.985 | 57630.856 | 57615.280 | 56822.557 |
| BIC | 57656.995 | 57659.370 | 57643.794 | 56851.072 |
| Log Likelihood | -28816.993 | -28812.428 | -28804.640 | -28408.279 |
| Deviance | 57633.985 | 57624.856 | 57609.280 | 56816.557 |
| Num. obs. | 99195 | 99195 | 99195 | 99195 |

\*\*\* $p < 0.001$ ; \*\* $p < 0.01$ ; \* $p < 0.05$

Table 5 focuses on topic and funding outcomes. Without consideration of other variables, an AAB preferred topic was associated with a lower probability of funding. However, after adjusting for IC award rate as well as other characteristics (whether the PI is an early stage investigator, whether the application included more than one PI, and whether the proposed research included human and/or animal subjects), there was no association between AAB preferred topics and funding.

**Table 5:** Probit Regression Models (regression coefficients and standard errors) with focus on topic type. Model 1 shows the univariable association of funding success according to whether the topic is AAB preferred. Model 2 shows results according to topic choice and IC award rate. Model 3 includes early stage investigator status, whether applications had multiple PIs, and whether the application included research on human subjects and/or animal subjects. AIC = Akaike Information Criterion; BIC = Bayesian Information Criterion; Num. obs. = Number of Observations. Other abbreviations same as in Tables 1 and 2.

|  | Model 1 | Model 2 | Model 3 |
| --- | --- | --- | --- |
| Intercept | -1.362***<br>(0.007) | -1.912***<br>(0.021) | -1.878***<br>(0.027) |
| AAB Preferred Topic | -0.044**<br>(0.013) | -0.006<br>(0.013) | -0.014<br>(0.015) |
| IC Award Rate |  | 0.062***<br>(0.002) | 0.058***<br>(0.002) |
| AAB Principal Investigator |  |  | -0.171***<br>(0.050) |
| Early Stage Investigator |  |  | 0.149***<br>(0.015) |
| Multi-PI Application |  |  | 0.100***<br>(0.014) |
| Human Subjects |  |  | -0.054***<br>(0.014) |
| Animal Subjects |  |  | -0.050***<br>(0.014) |
| AIC | 57642.323 | 56833.267 | 56693.521 |
| BIC | 57661.333 | 56861.781 | 56769.560 |
| Log Likelihood | -28819.162 | -28413.633 | -28338.760 |
| Deviance | 57638.323 | 56827.267 | 56677.521 |
| Num. obs. | 99195 | 99195 | 99195 |

\*\*\* $p < 0.001$ ; \*\* $p < 0.01$ ; \* $p < 0.05$

### Topics and Review Outcomes

To gain greater insights into possible peer review biases against topics preferred by AAB PIs, Figure 2, Panel A, shows the mean priority score by topic (of note, only discussed applications receive priority scores) according to the topic size, namely the number of submitted applications that were discussed for each topic. As would be expected topics of smaller size showed greater variability, a manifestation of regression to the mean.

**Figure 2:**
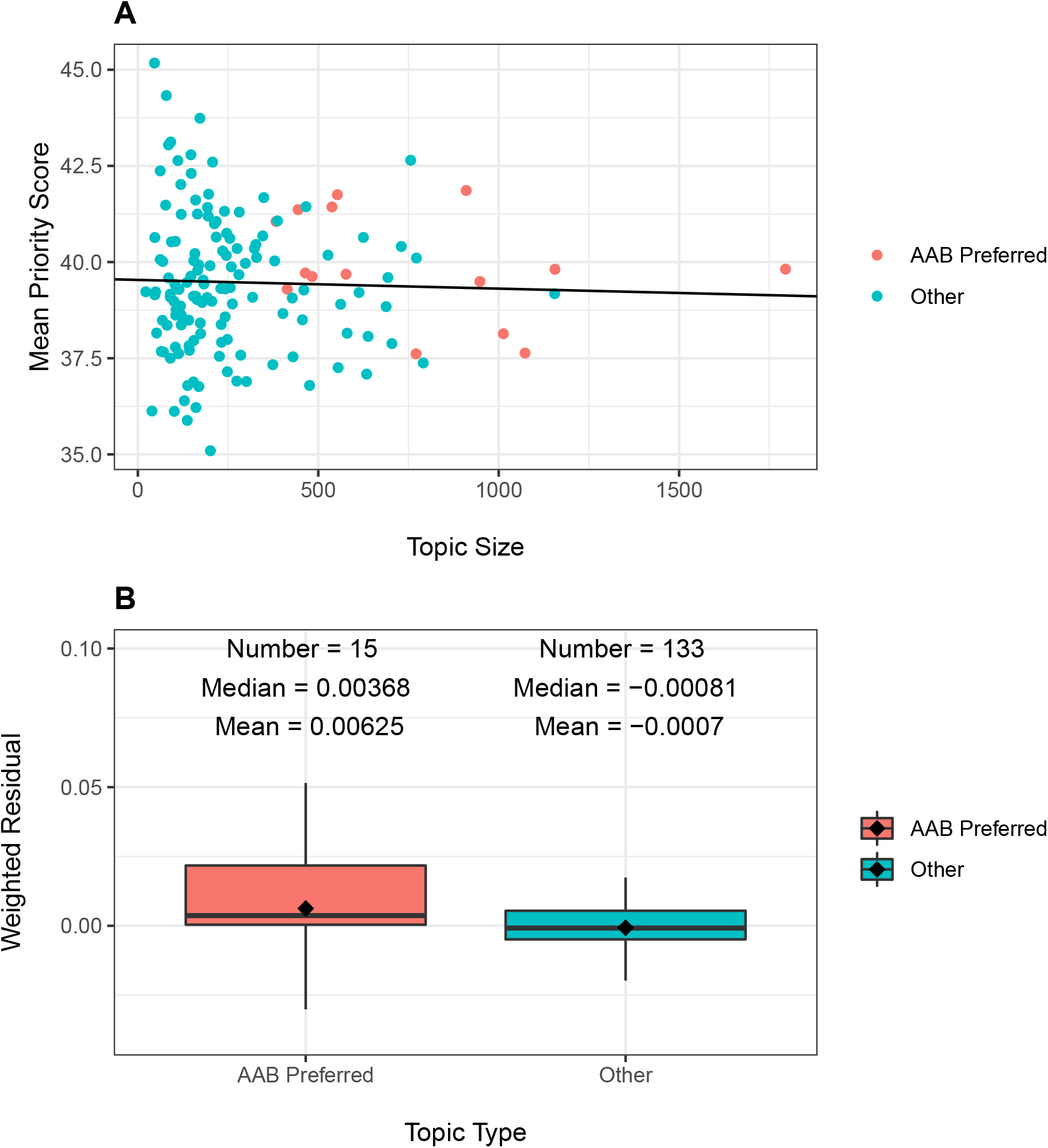
Topic peer review scores according to number of applications recevied (“Topic Size”) and topic type (AAB Preferred or Other). Panel A: Scatter plot of topic-specific mean peer review scores according to topic size. Each dot refers to a topic, with orange dots AAB preferred topics and green dots all others. The line is based on a linear regression of mean peer review scores on topic size. The slope of the line was not significant (P=0.68). Panel B: Distribution of weighted residuals of topic-specific mean review scores. Residuals are calculated as the distance between the dots in Panel A and the regression line, and are then weighted by topic size.

The line in Figure 2, Panel A, is based on a linear regression of predicted mean score according to topic size. Although the slope was slightly negative (coeffecient −0.0002264), the association was not significant (*p* = 0.68). Among AAB preferred topics (orange dots), there were 5 more than 1 point above the line (meaning with scores worse than predicted), while there were 3 more than 1 point below the line (meaning with scores better than predicted). The remaining 7 topics had mean scores that were within 1 point of the predicted value.

For each topic, we calculated a residual by subtracting from the topic-specific mean priority score the predicted mean priority score; we weighted the residuals by the topic size, as the larger topics contribute more information. Figure 2, Panel B, shows the distribution of the weighted residuals according to topic type. Residuals were more positive (i.e. worse) for AAB preferred topics. However, the absolute differences were small, much less than one priority score point (over a possible range of 10-90, with topic-specific mean values ranging from 35-45).

### Resubmission and Single-PI Applications

We repeated the main analyses, but this time focusing on resubmission applications. The findings were similar, except that, as expected, the absolute award rates were higher. We also conducted a separate series of analyses which repeated our primarily analyses but focusing solely on single-PI applications. Again, findings were similar. (See online supplements)

## Discussion

Among over 99,000 R01 applications submitted to NIH between 2011 and 2015, 2% were submitted by AAB PIs. Their applications were skewed towards a relatively small group of “AAB Preferred” topics; 10% of 148 topics accounted for 50% of AAB applications. Applications on AAB Preferred had similar review outcomes as those on other topics (Table 3) but were less likely to be funded. The lower award rates for AAB Preferred applications were associated assignment to ICs with lower overall award rates.

These observations reflect that there are two factors at play in determining whether an application submitted to NIH will be funded. The first, well known to all involved with NIH system, is peer review; those applications that receive better scores are more likely to be funded. But there is a second factor, namely the funding ecology of the IC to which the application is assigned. As shown in Table 2 applications with similar peer review outcomes are less likely to be funded if they are assigned to ICs with lower overall award rates. AAB PIs are more likely to submit applications to ICs with lower award rates, and applications (whether submitted by AAB or other PIs) that focus on AAB Preferred topics are more likely to be assigned to ICs with lower award rates (Figure 1).

Hoppe et al [Hoppe et al., 2019] found that topic choice partially accounted for funding disparities that adversely effect AAB PIs. We confirm this, but find that IC assignment (which, of course, is linked to topic) explains the disparities just as well, and that IC award rates explain the disparities even better (Table 4). Furthermore, after accounting for IC award rate, we found no association between topic choice and funding outcome (Table 5).

There is variability in how well different topics fare at peer review, but inspection of Figure 2 suggests that much of this variability reflects instability of estimates stemming from smaller sample sizes. Many topics that are not favored by AAB PIs receive better (lower) priority scores than the overall average, but many other such topics receive worse scores (Figure 2, Panel A). An inspection of weighted residuals suggest that AAB Preferred topics may fare a bit worse (Figure 2, Panel B), but to a much lower degree than the difference of award rates among assigned ICs (Table 1). Furthermore, it should be noted that applications on these topics were *more likely* to make it past the first hurdle of peer review, that is reaching the point of formal discussion (Table 3, see line “Discussed”); thus, if anything, peer reviewers may be slightly biased in favor of AAB-preferred topics.

Our primary analysis focused on first-time submissions of *de novo* applications in which the award rates were low (<10%). Nonetheless, our analyses of resubmission applications yielded similar findings (online Supplement). It is also important to note that this was an analysis of applications, not persons. This is an issue when considering multi-PI applications, since all PI’s, not just the contact PI, plays a role in choosing the topic of their proposal. We effectively treat the contact PI as “above among equals.” Nonetheless, an analysis confined to single PI applications yielded similar findings (online Supplement).

## Conclusion

The lower rate of funding for applications focused on AAB Preferred topics is likely primarily due to their assignment to ICs with lower award rates. These applications have similar peer review outcomes as those focused on other topics. Topic choice does partially explain race-based funding disparities, but IC-specific award rates explain the disparities to an even greater degree. After accounting for IC-specific award rates, we find no association between topic choice and funding outcomes.

## Supplementary Materials: Resubmission Applications

In our primary analyses, we excluded resubmission applications and competing renewals. Since many applications are not funded on the first try, we repeated our analyses focusing on resubmissions. Our findings are similar.

Of 32,518 resubmission applications received, 9605 were funded, for an overall award rate of 30%. There were 451 applications, or 1%, submitted by AAB PIs.

Table 1 shows review and funding outcomes for applications according to whether the assignment was to an IC in the top quartile of AAB resubmission applications (“Higher AAB”). These ICs were the National Institute of Dental and Craniofacial Research, the National Institute of Environmental Health Sciences, the National Institute of Child Health and Human Development, the National Institute of Minority Health and Disparities, the National Institute of Nursing Research, and the Fogarty International Center. Applications submitted to Higher AAB ICs were 3 times more likely to come from AAB PIs. Review outcomes – proportion discussed and, for those applications that were discussed at peer review meetings, priority scores and percentile rankings – were similar in both groups. Despite the similar review outcomes, they were 16% less likely to be funded.

Table 2 shows corresponding values according to whether applications were focused on the 16 topics that made up 50% of all resubmission applications with AAB PIs (“AAB Preferred” topics). Again, peer review outcomes were similar in the two groups, but applications focusing on AAB Preferred topics were 3% less likely to be funded.

Table 3 shows the association of an application with an AAB PI with the probability of funding. Consistent with Hoppe et al [1] and prior literature [2], AAB PI applications had a lower likelihood of funding (negative regression coefficient for AAB Principal Investigator). Adjusting for the topic (AAB Preferred or Other) reduced the regression coefficient for race by 3%; adjusting for IC assignment (Higher or Lower AAB) reduced the regression coefficient by 11%. Adjusting for the award rate of the assigned IC reduced the regression coefficient for race by 10%.

Table 4 focuses on topic and funding outcomes. Without consideration of other variables, an AAB preferred topic was associated with a lower probability of funding. However, after adjusting for IC award rate as well as other characteristics (whether the PI is an early stage investigator, whether the application included more than one PI, and whether the proposed research included human and/or animal subjects), there was no association between AAB preferred topics and funding.

**Table 1:** Resubmission application review and funding outcomes according to whether Institute or Center received a higher or lower proportion of applications from AAB principal investigators. AAB = African American or Black; PI = Principal Investigator.

| Characteristic or Outcome |  | Higher AAB | Lower AAB |
| --- | --- | --- | --- |
| Total N (%) |  | 3300 (10.1) | 29218 (89.9) |
| PI AAB | Yes | 103 (3.1) | 348 (1.2) |
| Discussed | Yes | 2382 (72.2) | 21025 (72.0) |
| Priority Score | Median (IQR) | 31.0 (21.0 to 41.0) | 31.0 (22.0 to 40.0) |
| Percentile Ranking | Median (IQR) | 21.0 (9.0 to 36.0) | 21.0 (10.0 to 35.0) |
| Awarded | Yes | 836 (25.3) | 8769 (30.0) |

**Table 2:** Resubmission application review and funding outcomes according to whether topic was among those that accounted for the majority of AAB applications. Abbreviations as in Table 2.

| Characteristic or Outcome |  | AAB Preferred | Other |
| --- | --- | --- | --- |
| Total N (%) |  | 8356 (25.7) | 24162 (74.3) |
| PI AAB | Yes | 224 (2.7) | 227 (0.9) |
| Discussed | Yes | 6107 (73.1) | 17300 (71.6) |
| Priority Score | Median (IQR) | 31.0 (22.0 to 40.0) | 31.0 (22.0 to 40.0) |
| Percentile Ranking | Median (IQR) | 21.0 (9.8 to 36.0) | 21.0 (10.0 to 35.0) |
| Assigned IC AAB Proportion | Higher AAB | 1491 (17.8) | 1809 (7.5) |
| Awarded | Yes | 2419 (28.9) | 7186 (29.7) |

To gain greater insights into possible peer review biases against topics preferred by AAB PIs, Figure 1, Panel A, shows the mean priority score by topic (of note, only discussed applications receive priority scores) according to the topic size, namely the number of submitted applications that were discussed for each topic. As would be expected topics of smaller size showed greater variability, a manifestation of regression to the mean.

The line in Figure 1, Panel A, is based on a linear regression of predicted mean score according to topic size. Although the slope was slightly negative (coeffecient −0.0017232), the association was not significant (*p* = 0.14). Among AAB preferred topics (orange dots), there were 4 more than 1 point above the line (meaning with scores worse than predicted), while there were 6 more than 1 point below the line (meaning with scores better than predicted). The remaining 6 topics had mean scores that were within 1 point of the predicted value.

For each topic, we calculated a residual by subtracting from the topic-specific mean priority score the predicted mean priority score; we weighted the residuals by the topic size, as the larger topics contribute more information. Figure 1, Panel B, shows the distribution of the weighted residuals according to topic type. Residuals were more positive (i.e. worse) for AAB preferred topics. However, the absolute differences were small, much less than one priority score point (over a possible range of 10-90, with topic-specific mean values ranging from 35-45).

**Table 3:** Probit Regression Models (regression coefficients and standard errors) of resubmission applications with focus on the PI. Model 1 shows the univariable association of funding success according to whether the PI is AAB. Model 2 adjusts for topic, Model 3 adjusts for IC assignment, and Model 4 adjusts for IC award rate. Note that the absolute value for the regression coefficient linking AAB PI to funding outcome decreases with each of these adjustments, with the greatest reduction after adjusting for IC award rate. AIC = Akaike Information Criterion; BIC = Bayesian Information Criterion; Num. obs. = Number of Observations. Other abbreviations as in Tables 1 and 2.

|  | Model 1 | Model 2 | Model 3 | Model 4 |
| --- | --- | --- | --- | --- |
| Intercept | −0.536***<br>(0.007) | −0.531***<br>(0.009) | −0.522***<br>(0.008) | −1.400***<br>(0.065) |
| AAB Principal Investigator | −0.158*<br>(0.065) | −0.153*<br>(0.065) | −0.141*<br>(0.065) | −0.142*<br>(0.065) |
| AAB Preferred Topic |  | −0.020<br>(0.017) |  |  |
| IC Higher AAB Applications |  |  | −0.137***<br>(0.025) |  |
| IC Award Rate |  |  |  | 0.029***<br>(0.002) |
| AIC | 39467.941 | 39468.474 | 39439.201 | 39287.118 |
| BIC | 39484.721 | 39493.643 | 39464.369 | 39312.287 |
| Log Likelihood | −19731.971 | −19731.237 | −19716.600 | −19640.559 |
| Deviance | 39463.941 | 39462.474 | 39433.201 | 39281.118 |
| Num. obs. | 32518 | 32518 | 32518 | 32518 |

\*\*\* $p < 0.001$ ; \*\* $p < 0.01$ ; \* $p < 0.05$

**Table 4:** Probit Regression Models (regression coefficients and standard errors) of resubmission applications with focus on topic type. Model 1 shows the univariable association of funding success according to whether the topic is AAB preferred. Model 2 shows results according to topic choice and IC award rate. Model 3 includes early stage investigator status, whether applications had multiple PIs, and whether the application included research on human subjects and/or animal subjects. AIC = Akaike Information Criterion; BIC = Bayesian Information Criterion; Num. obs. = Number of Observations. Other abbreviations as in Tables 1 and 2.

|  | Model 1 | Model 2 | Model 3 |
| --- | --- | --- | --- |
| Intercept | −0.532***<br>(0.008) | −1.404***<br>(0.065) | −1.429***<br>(0.068) |
| AAB Preferred Topic | −0.023<br>(0.017) | −0.001<br>(0.017) | −0.005<br>(0.018) |
| IC Award Rate |  | 0.029***<br>(0.002) | 0.029***<br>(0.002) |
| AAB Principal Investigator |  |  | −0.159*<br>(0.065) |
| Early Stage Investigator |  |  | 0.218***<br>(0.019) |
| Multi-PI Application |  |  | 0.036<br>(0.020) |
| Human Subjects |  |  | −0.018<br>(0.018) |
| Animal Subjects |  |  | −0.007<br>(0.018) |
| AIC | 39472.107 | 39291.937 | 39165.838 |
| BIC | 39488.886 | 39317.106 | 39232.955 |
| Log Likelihood | −19734.053 | −19642.968 | −19574.919 |
| Deviance | 39468.107 | 39285.937 | 39149.838 |
| Num. obs. | 32518 | 32518 | 32518 |

\*\*\* $p < 0.001$ ; \*\* $p < 0.01$ ; \* $p < 0.05$

**Figure 1:**
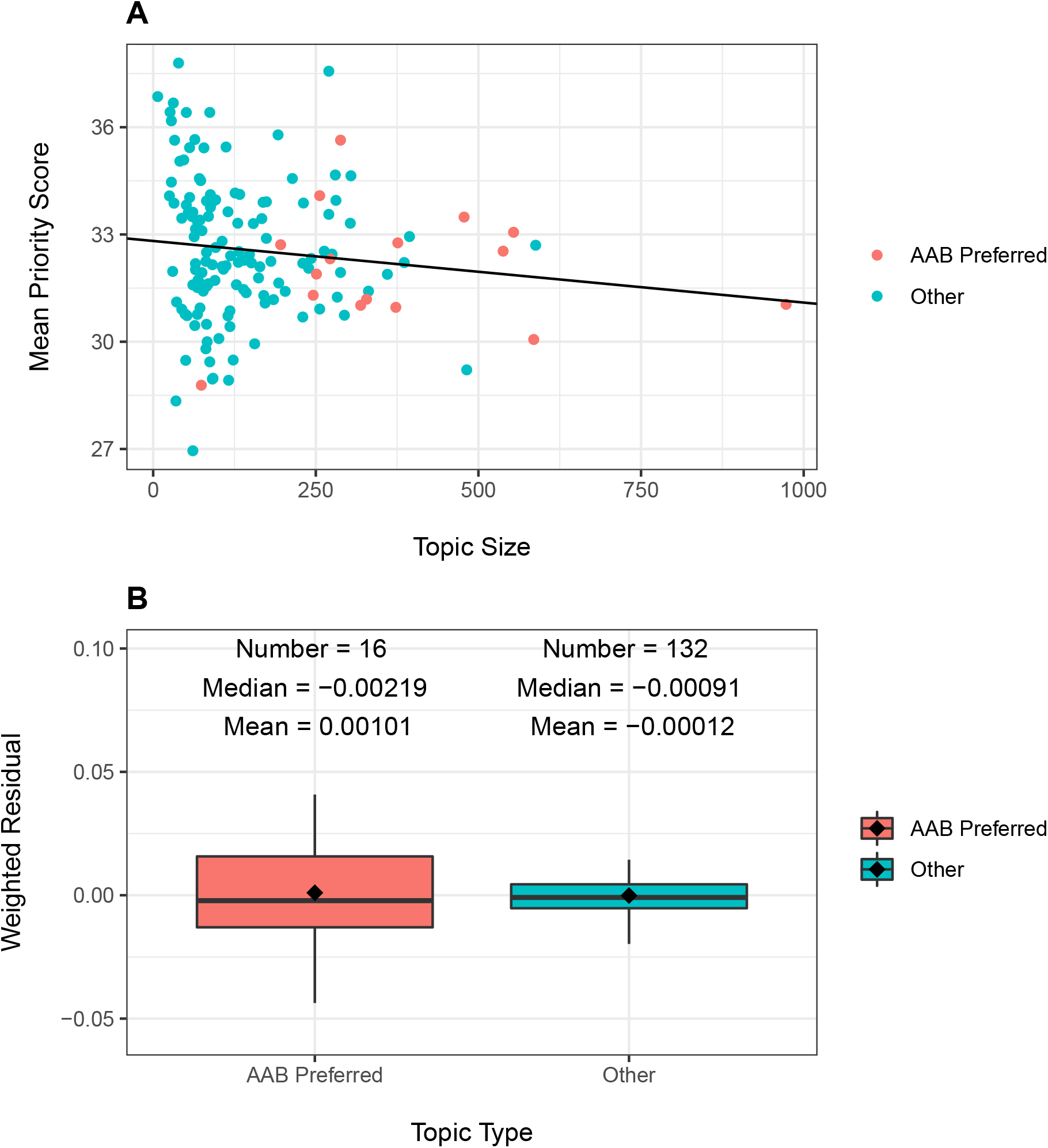
Topic peer review scores according to number of resubmission applications recevied (“Topic Size”) and topic type (AAB Preferred or Other). Panel A: Scatter plot of topic-specific mean peer review scores according to topic size. Each dot refers to a topic, with orange dots AAB preferred topics and green dots all others. The line is based on a linear regression of mean peer review scores on topic size. The slope of the line was not significant (P=0.14). Panel B: Distribution of weighted residuals of topic-specific mean review scores. Residuals are calculated as the distance between the dots in Panel A and the regression line, and are then weighted by topic size.

## Supplementary Materials: Single PI Applications

Here we repeat our primary analyses, but focus on single PI applications.

Of 79,016 *de novo* first-time single-PI applications received, 6590 were funded, for an overall award rate of 8%. There were 1357 applications, or 2%, submitted by AAB PIs.

Table 1 shows review and funding outcomes for applications according to whether the assignment was to an IC in the top quartile of AAB single-PI applications (“Higher AAB”). Applications submitted to Higher AAB ICs were 3 times more likely to come from AAB PIs. They were slightly less likely to be discussed (44% vs 45%), but for those applications that were discussed at peer review meetings, priority scores and percentile rankings were similar in both groups. Despite the similar review outcomes, they were 21% less likely to be funded.

Table 2 shows corresponding values according to whether applications were focused on the 16 topics that made up 50% of all single-PI applications with AAB PIs (“AAB Preferred” topics). Applications on AAB Preferred topics were slightly more likely to be discussed (46% vs 44%) while other peer review outcomes were similar in the two groups. Applications focusing on AAB Preferred topics were 7% less likely to be funded.

Table 3 shows the association of an application with an AAB PI with the probability of funding. Consistent with Hoppe et al [1] and prior literature [2], AAB PI applications had a lower likelihood of funding (negative regression coefficient for AAB Principal Investigator). Adjusting for the topic (AAB Preferred or Other) reduced the regression coefficient for race by 4%; adjusting for IC assignment (Higher or Lower AAB) reduced the regression coefficient by 9%. Adjusting for the award rate of the assigned IC reduced the regression coefficient for race by 15%.

Table 4 focuses on topic and funding outcomes. Without consideration of other variables, an AAB preferred topic was associated with a lower probability of funding. However, after adjusting for IC award rate as well as other characteristics (whether the PI is an early stage investigator, and whether the proposed research included human and/or animal subjects), there was no association between AAB preferred topics and funding.

**Table 1:** Single-PI application review and funding outcomes according to whether Institute or Center received a higher or lower proportion of applications from AAB principal investigators. AAB = African American or Black; PI = Principal Investigator.

| Characteristic or Outcome |  | Higher AAB | Lower AAB |
| --- | --- | --- | --- |
| Total N (%) |  | 15982 (20.2) | 63034 (79.8) |
| PI AAB | Yes | 505 (3.2) | 852 (1.4) |
| Discussed | Yes | 6988 (43.7) | 28384 (45.0) |
| Priority Score | Median (IQR) | 39.0 (30.0 to 48.0) | 40.0 (31.0 to 48.0) |
| Percentile Ranking | Median (IQR) | 34.0 (21.0 to 45.0) | 34.0 (21.0 to 45.0) |
| Awarded | Yes | 1109 (6.9) | 5481 (8.7) |

**Table 2:**
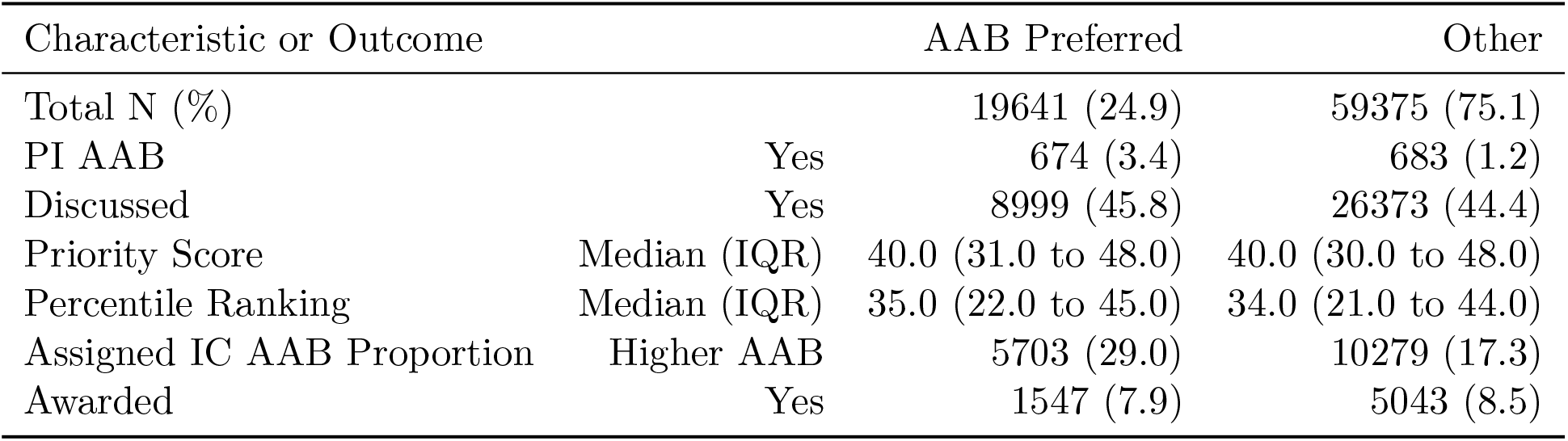
Single-PI application review and funding outcomes according to whether topic was among those that accounted for the majority of AAB applications. Abbreviations as in Table 2.

The line in Figure 1, Panel A, is based on a linear regression of predicted mean score according to topic size. Although the slope was slightly negative (coeffecient −0.0005473), the association was not significant (*p* = 0.46). Among AAB preferred topics (orange dots), there were 7 more than 1 point above the line (meaning with scores worse than predicted), while there were 4 more than 1 point below the line (meaning with scores better than predicted). The remaining 5 topics had mean scores that were within 1 point of the predicted value.

**Table 3:** Probit Regression Models (regression coefficients and standard errors) of single-PI applications with focus on the PI. Model 1 shows the univariable association of funding success according to whether the PI is AAB. Model 2 adjusts for topic, Model 3 adjusts for IC assignment, and Model 4 adjusts for IC award rate. Note that the absolute value for the regression coefficient linking AAB PI to funding outcome decreases with each of these adjustments, with the greatest reduction after adjusting for IC award rate. AIC = Akaike Information Criterion; BIC = Bayesian Information Criterion; Num. obs. = Number of Observations. Other abbreviations as in Tables 1 and 2.

|  | Model 1 | Model 2 | Model 3 | Model 4 |
| --- | --- | --- | --- | --- |
| Intercept | −1.379***<br>(0.006) | −1.370***<br>(0.007) | −1.357***<br>(0.007) | −1.906***<br>(0.022) |
| AAB Principal Investigator | −0.223***<br>(0.056) | −0.214***<br>(0.056) | −0.203***<br>(0.056) | −0.189***<br>(0.057) |
| AAB Preferred Topic |  | −0.037*<br>(0.015) |  |  |
| IC Higher AAB Applications |  |  | −0.117***<br>(0.017) |  |
| IC Award Rate |  |  |  | 0.061***<br>(0.002) |
| AIC | 45341.889 | 45337.935 | 45293.374 | 44665.262 |
| BIC | 45360.443 | 45365.767 | 45321.206 | 44693.094 |
| Log Likelihood | −22668.944 | −22665.967 | −22643.687 | −22329.631 |
| Deviance | 45337.889 | 45331.935 | 45287.374 | 44659.262 |
| Num. obs. | 79016 | 79016 | 79016 | 79016 |

\*\*\* $p < 0.001$ ; \*\* $p < 0.01$ ; \* $p < 0.05$

**Table 4:** Probit Regression Models (regression coefficients and standard errors) of single-PI applications with focus on topic type. Model 1 shows the univariable association of funding success according to whether the topic is AAB preferred. Model 2 shows results according to topic choice and IC award rate. Model 3 includes early stage investigator status, whether applications had multiple PIs, and whether the application included research on human subjects and/or animal subjects. AIC = Akaike Information Criterion; BIC = Bayesian Information Criterion; Num. obs. = Number of Observations. Other abbreviations as in Tables 1 and 2.

|  | Model 1 | Model 2 | Model 3 |
| --- | --- | --- | --- |
| Intercept | -1.373***<br>(0.007) | -1.910***<br>(0.022) | -1.866***<br>(0.030) |
| AAB Preferred Topic | -0.041**<br>(0.015) | 0.000<br>(0.015) | 0.006<br>(0.016) |
| IC Award Rate |  | 0.061***<br>(0.002) | 0.057***<br>(0.002) |
| AAB Principal Investigator |  |  | -0.207***<br>(0.057) |
| Early Stage Investigator |  |  | 0.160***<br>(0.015) |
| Human Subjects |  |  | -0.068***<br>(0.016) |
| Animal Subjects |  |  | -0.035*<br>(0.015) |
| AIC | 45351.399 | 44677.087 | 44550.569 |
| BIC | 45369.954 | 44704.920 | 44615.511 |
| Log Likelihood | -22673.700 | -22335.544 | -22268.284 |
| Deviance | 45347.399 | 44671.087 | 44536.569 |
| Num. obs. | 79016 | 79016 | 79016 |

\*\*\* $p < 0.001$ ; \*\* $p < 0.01$ ; \* $p < 0.05$

**Figure 1:**
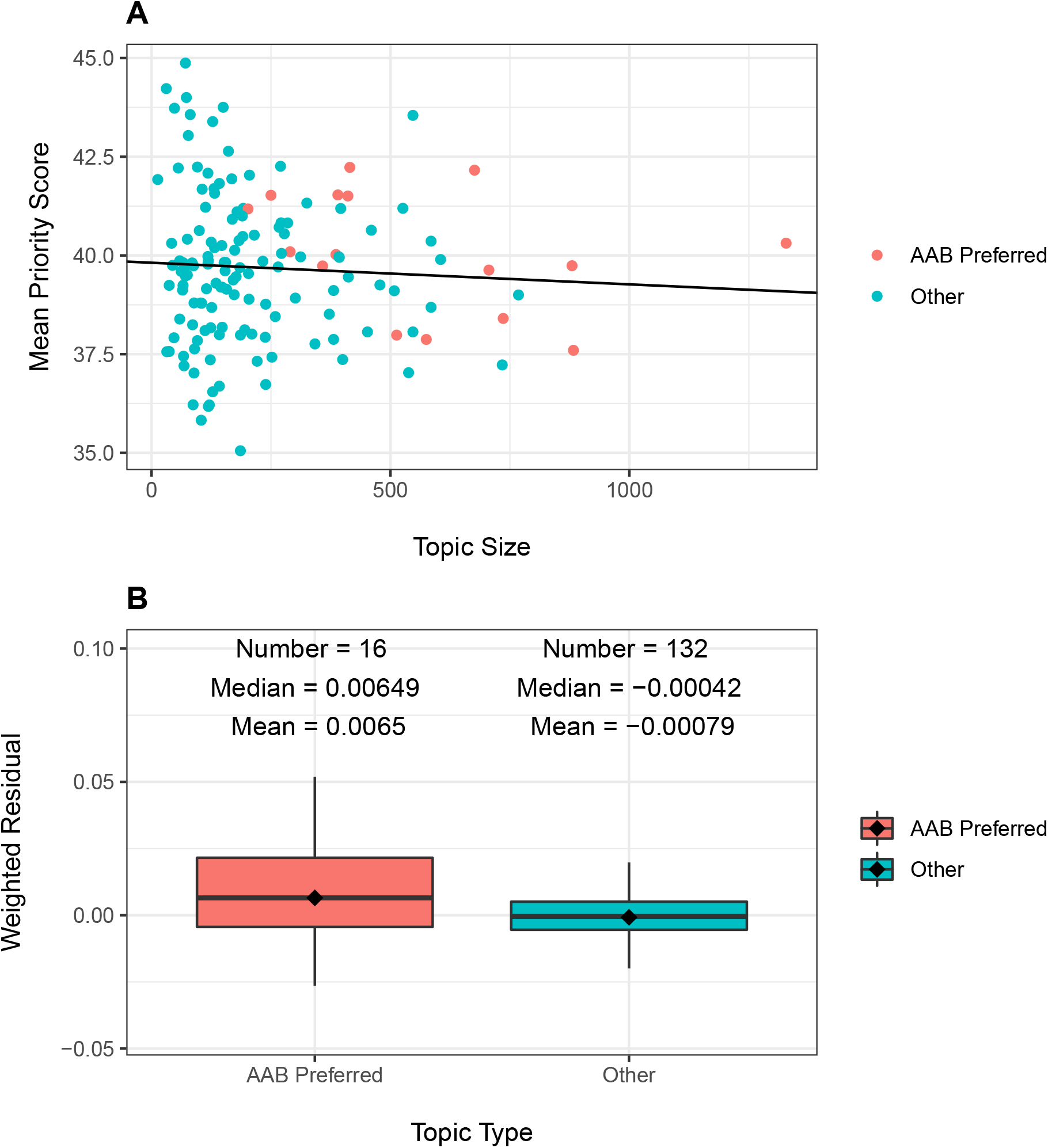
Topic peer review scores according to number of single-PI applications recevied (“Topic Size”) and topic type (AAB Preferred or Other). Panel A: Scatter plot of topic-specific mean peer review scores according to topic size. Each dot refers to a topic, with orange dots AAB preferred topics and green dots all others. The line is based on a linear regression of mean peer review scores on topic size. The slope of the line was not significant (P=0.46). Panel B: Distribution of weighted residuals of topic-specific mean review scores. Residuals are calculated as the distance between the dots in Panel A and the regression line, and are then weighted by topic size.

